# Simphony: simulating large-scale, rhythmic data

**DOI:** 10.1101/497859

**Authors:** Jordan M. Singer, Darwin Y. Fu, Jacob J. Hughey

**Author notes:** To whom all correspondence should be addressed.

## Abstract

Simulated data are invaluable for assessing a computational method’s ability to distinguish signal from noise. Although many biological systems show rhythmicity, there is no general-purpose tool to simulate large-scale, rhythmic data. Here we present Simphony, an R package for simulating data from experiments in which the abundances of rhythmic and non-rhythmic features (e.g., genes) are measured at multiple time points in multiple conditions. Simphony has parameters for specifying experimental design and each feature’s rhythmic properties (e.g., shape, amplitude, and phase). In addition, Simphony can sample measurements from Gaussian and negative binomial distributions, the latter of which approximates read counts from next-generation sequencing data. We show an example of using Simphony to benchmark a method for detecting rhythms. Our results suggest that Simphony can aid experimental design and computational method development. Simphony is thoroughly documented and freely available at https://github.com/hugheylab/simphony.

## Introduction

Rhythms are ubiquitous across domains of life and across timescales, from hourly division of bacteria (Cooper & Helmstetter, 1968) to seasonal growth of trees (Kramer, 1936). These biological rhythms are often driven by systems of genes and proteins. Prominent examples are the systems underlying circadian rhythms, which have a period of approximately 24 hours and have been observed in species across the biosphere (Young & Kay, 2001).

To interrogate these rhythmic biological systems, researchers are increasingly using technologies that measure the abundance of thousands of molecules in parallel, e.g., the transcriptome or proteome. The critical decisions then become how to design the experiments and how to analyze the data. For example, there are now numerous methods for detecting rhythms in high-dimensional data (Wu et al., 2016). A valuable aid to such decisions is simulation. In simulated data, unlike in experimental data, the ground truth (e.g., whether a gene is rhythmic) is known. Consequently, as long as the simulated data recapitulate the essential features of experimental data, they enable one to fairly estimate a method’s performance in a given experimental design (Love, Hogenesch & Irizarry, 2016). Simulated data are also faster and less expensive to generate than experimental data, especially omics data from high-resolution time courses.

Unfortunately, there is a shortage of publicly available tools for simulating rhythmic data. This forces researchers (both data generators and method developers) to create their own simulation framework from scratch or to forgo simulations altogether. Recently, a tool called CircaInSilico was developed to begin to fill this gap (Hughes et al., 2017). Although CircaInSilico has a convenient user interface, it has several limitations. For example, the simulated rhythms can only be sinusoidal. In addition, even though read counts from next-generation sequencing data are often modeled using a negative binomial distribution (Robinson & Smyth, 2007), CircaInSilico can only simulate Gaussian noise. A more general tool for simulating RNA-seq reads is Polyester (Frazee et al., 2015). Although Polyester can simulate reads from multiple conditions or time points, it is not specifically designed to simulate rhythms. Furthermore, Polyester models many aspects of the sequencing process, which introduces considerable computational costs and may not be directly relevant for designing experiments to collect rhythmic data or evaluating methods to analyze such data. Thus, there is still a need for a flexible, easy-to-use tool to simulate large-scale, rhythmic data.

To address this need, we developed a simulation package called Simphony. Simphony has adjustable parameters for specifying experimental design and modeling rhythms, including the ability to sample from Gaussian and negative binomial distributions. Simphony is implemented in R, thoroughly documented, and freely available at https://github.com/hugheylab/simphony.

## Materials and Methods

All data and code to reproduce this study are available at https://figshare.com/s/549da44928b243df47d5.

### Simulating rhythmic data using Simphony

Simphony simulates experiments in which the abundances of rhythmic and non-rhythmic features (e.g., genes) are measured at multiple time points in one or more conditions. Within a given simulated experiment (i.e., a simulation), the expected abundance *m* of feature *i* in condition *k* at time *t* is modeled as

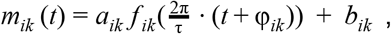

where *a* is the amplitude, *f* is a periodic function with period 2*π* (by default, *f* (θ) = *sin*(θ)), τ is the period of rhythmic changes in abundance (by default, 24), φ is the phase, and *b* is the baseline abundance. Non-rhythmicity is defined by *a* = 0. Given *m_ik_*(*t*), Simphony samples measurements from one of two families of distributions: Gaussian and negative binomial. The former represents an idealized experimental scenario, whereas the latter approximates read counts from next-generation sequencing. For Gaussian sampling, the abundance of feature *i* in sample *j* belonging to condition *k* follows

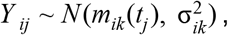

where σ^2^ is the variance (by default, 1). For negative binomial sampling, we follow a similar strategy to DESeq2 (Love, Huber & Anders, 2014) and Polyester (Frazee et al., 2015), such that

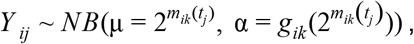

where μ is the expected counts, *α* is the dispersion (the variance of a negative binomial distribution is *V ar*(*Y*) = μ + αμ^2^), and *g* is a function that maps expected counts to dispersion. The default *g* was estimated from RNA-seq data from mouse liver (see the next section for details).

Experimental design in Simphony is specified in one of three ways: (1) interval between time points and number of samples per time point per condition, (2) exact time points and number of samples per time point per condition, or (3) time points from a uniform distribution (between 0 and τ) and total number of samples per condition.

The Simphony R package has three dependencies: data.table (Dowle & Srinivasan, 2018), foreach (Microsoft & Weston, 2017), and locfit (Loader, 2013).

### Estimating statistical properties of experimental RNA-seq data

To estimate the relationship between expected counts and dispersion in real RNA-seq data, we used PRJNA297287 (Atger et al., 2015). We used the samples that were collected in quadruplicate from livers of wild-type, ad libitum-fed mice every 2 hours for 24 hours in LD 12:12 (48 samples total). We downloaded the raw reads, then quantified gene-level counts using Salmon v0.11.3 (Patro et al., 2017) and tximport v1.8.0 (Soneson, Love & Robinson, 2015). We kept the 15,069 genes that had at least 10 counts in half of the samples. We used DESeq2 v1.20.0 to estimate parametric and local regression-based mean-dispersion curves (Love, Huber & Anders, 2014) (Fig. S1A). The input to DESeq2 included a design matrix based on cosinor regression, so that dispersion estimates were not biased by variation in expression due to a daily rhythm. Compared to the parametric mean-dispersion curve, the local regression-based curve had a considerably lower root-mean-squared error (0.94 compared to 1.09, in units of log dispersion), so we set it as the default in Simphony. DESeq2 also provided an estimate of the variance of the residual log dispersion (around the curve). Finally, we used fitdistrplus (Delignette-Muller & Dutang, 2015) to approximate the distribution of mean normalized counts as log-normal. The Simphony documentation includes an example of how to sample from the estimated distributions of residual log dispersion and mean normalized counts (Fig. S1B).

### Validating statistical properties of simulated data

We performed multiple simulations to validate the statistical properties of data generated by Simphony. Each simulation had time points spaced 0.1 h apart (period of 24 h), 100 samples per time point, and one gene for each unique combination of parameter values related to gene expression. Simulations based on negative binomial sampling used the default function for calculating dispersion.

To validate mean and standard deviation, we simulated non-rhythmic expression (amplitude of 0) based on Gaussian and negative binomial sampling. For the simulation using Gaussian sampling, we varied the desired mean and standard deviation. For the simulation using negative binomial sampling, we varied the desired mean log_2_ counts. In both cases, we then calculated the empirical mean and standard deviation (Table S1).

To validate amplitude and phase, we simulated rhythmic expression based on Gaussian and negative binomial sampling (using the default *f* (θ) = *sin*(θ)). For both types of sampling, we varied the desired amplitude and phase. For the simulation based on Gaussian sampling, we used the limma R package v3.36.5 (Smyth, 2004; Ritchie et al., 2015) to fit each gene’s expression to a linear model that had terms for 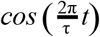 and 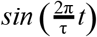 (cosinor regression). We then used the model coefficients to estimate each gene’s amplitude and phase according to the trigonometric identity *a* · *cos*θ + *b* · *sin*θ = *c* · *sin*(θ + φ), where 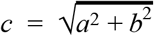 and 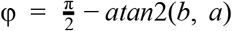 (Table S2). For the simulation based on negative binomial sampling, we followed a similar procedure, except we log-transformed the expression counts before passing them to limma.

### Detecting rhythmicity in simulated data

We used limma to calculate gene-wise p-values of rhythmicity. The p-value was based on a moderated F-test on the coefficients corresponding to the variables 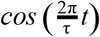 and 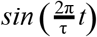 (cosinor regression). Because the expression values were counts sampled from the negative binomial family, we first transformed them using log_2_(counts+1). We used limma with the default settings, except we allowed it to fit a mean-variance trend across genes (limma-trend) (Law et al., 2014). We then used the p-values and the precrec R package v0.9.1 (Saito & Rehmsmeier, 2017) to calculate the area under the receiver operating characteristic (ROC) curve for distinguishing non-rhythmic genes from each group of rhythmic genes (specified by rhythm amplitude and baseline in log_2_ counts).

## Results

To validate the statistical properties of data generated by Simphony, we simulated data covering a range of parameter values for the Gaussian and negative binomial families. To ensure that the properties approached their asymptotic values, time points were spaced 0.1 h apart (period of 24 h), each with 100 samples. For non-rhythmic expression, we verified that the observed mean and standard deviation corresponded to the expected values (Table S1). For rhythmic expression, we verified that the observed amplitude and phase corresponded to the expected values (Materials and Methods; Table S2).

To highlight Simphony’s flexibility, we simulated experiments in which rhythmic gene expression followed a sinusoid or a sawtooth wave, with expression values sampled from the Gaussian or negative binomial family (Fig. 1). We also simulated an experiment having two conditions, in which genes’ rhythms had a different amplitude or phase in each condition (Fig. S2).

**Figure 1.**
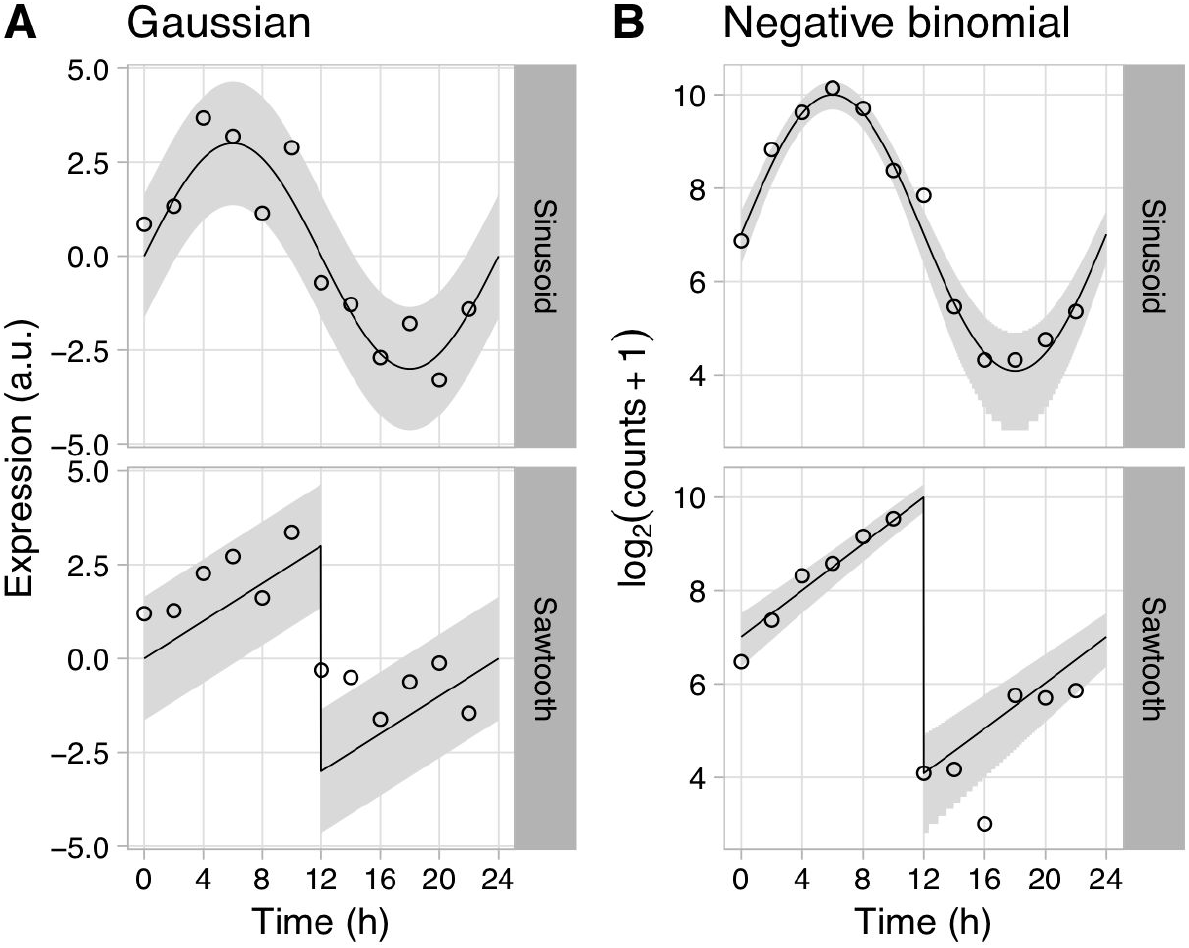
Examples of rhythmic data generated by Simphony. Gene expression values were sampled from the **(A)** Gaussian or **(B)** negative binomial family. Rhythms followed a sinusoid or a sawtooth wave of period 24 h and amplitude 4. For Gaussian sampling, the baseline expression was 0 and the standard deviation was 1. For negative binomial sampling, the baseline log_2_ counts was 7. Time points were spaced 2 h apart, with 1 sample per time point. Circles show the sampled gene expression values, black lines show the expected expression over time, and gray ribbons show the corresponding 90% prediction intervals. The prediction intervals for negative binomial sampling have discontinuities because the sampled values can only be integers greater than or equal to zero.

To illustrate Simphony’s utility, we used it to benchmark cosinor regression, a method for detecting rhythms (Halberg, Tong & Johnson, 1967). We simulated experiments having various intervals between time points and one sample per time point. Each simulation included 20,000 genes spanning a range of values for baseline expression and rhythm amplitude (including amplitude 0 for non-rhythmic genes) (Fig. S3A). For each simulation, we calculated each gene’s p-value of rhythmicity using limma (Materials and Methods), then calculated the area under the ROC curve for distinguishing non-rhythmic genes from each group of rhythmic genes. As expected, rhythm detection improved as rhythm amplitude increased or the interval between time points decreased (Fig. 2A). Rhythm detection also improved as baseline expression increased (and thus as the standard deviation of log-transformed counts of non-rhythmic genes decreased; Fig. 2B and Fig. S3B).

**Figure 2.**
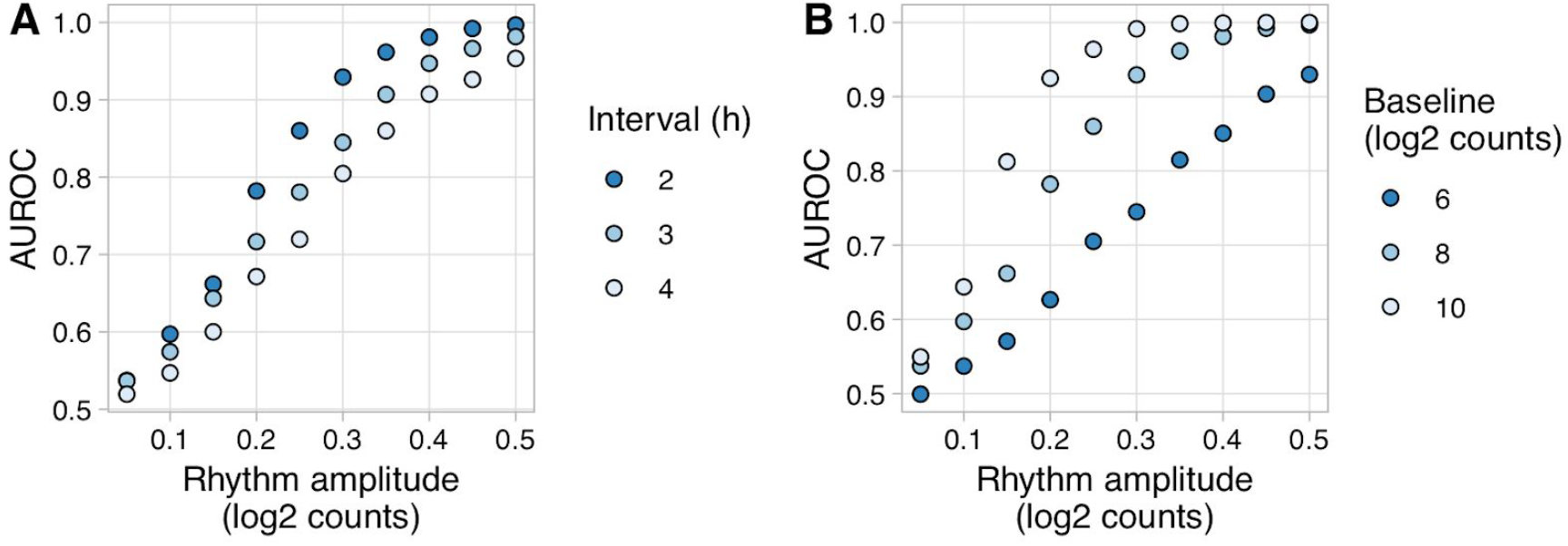
Evaluating rhythm detection using data generated by Simphony. Simulations had various values of the interval between time points and one replicate per time point. Each simulation included 20,000 genes having various values of baseline expression and rhythm amplitude (including amplitude 0). Rhythms followed a sinusoid of period 24 h. Expression values were sampled from the negative binomial family. We used the gene-wise p-values of rhythmicity (calculated using limma) to calculate the area under the ROC curve (AUROC) for distinguishing non-rhythmic genes from each group of rhythmic genes. **(A)** AUROC vs. rhythm amplitude and interval, for genes with a baseline log_2_ counts of 8. **(B)** AUROC vs. rhythm amplitude and baseline expression, for the simulation with an interval of 2 h. AUROC of 0.5 corresponds to random detection, while AUROC of 1 corresponds to perfect detection.

## Conclusions

Simphony is a versatile framework for simulating rhythmic data. Although Simphony is especially apt for simulating circadian transcriptome data, it is general enough to simulate data of various types (e.g., bioluminescence). In the future, we plan to extend Simphony to simulate non-stationary trends (e.g., damped rhythms) and to accommodate different periods for different genes within a simulation. Altogether, Simphony can guide the design of experiments for interrogating rhythmic biological systems and help benchmark methods for analyzing data containing rhythmic signals.

## References

Atger F., Gobet C., Marquis J., Martin E., Wang J., Weger B., Lefebvre G., Descombes P., Naef F., Gachon F. 2015. Circadian and feeding rhythms differentially affect rhythmic mRNA transcription and translation in mouse liver. Proceedings of the National Academy of Sciences of the United States of America 112:E6579–88.

Cooper S., Helmstetter CE. 1968. Chromosome replication and the division cycle of Escherichia coli B/r. Journal of molecular biology 31:519–540.

Delignette-Muller M., Dutang C. 2015. fitdistrplus: An R Package for Fitting Distributions. Journal of Statistical Software, Articles 64:1–34.

Dowle M., Srinivasan A. 2018. data.table: Extension of ‘data.frame’. Comprehensive R Archive Network (CRAN).

Frazee AC., Jaffe AE., Langmead B., Leek JT. 2015. Polyester: simulating RNA-seq datasets with differential transcript expression. Bioinformatics 31:2778–2784.

Halberg F., Tong YL., Johnson EA. 1967. Circadian System Phase — An Aspect of Temporal Morphology; Procedures and Illustrative Examples. In: von Mayersbach H ed. The Cellular Aspects of Biorhythms: Symposium on Rhythmic Research Sponsored by the VIIIth International Congress of Anatomy Wiesbaden 8.–14. August 1965. Berlin, Heidelberg: Springer Berlin Heidelberg, 20–48.

Hughes ME., Abruzzi KC., Allada R., Anafi R., Arpat AB., Asher G., Baldi P., de Bekker C., Bell-Pedersen D., Blau J., Brown S., Ceriani MF., Chen Z., Chiu JC., Cox J., Crowell AM., DeBruyne JP., Dijk D-J., DiTacchio L., Doyle FJ., Duffield GE., Dunlap JC., Eckel-Mahan K., Esser KA., FitzGerald GA., Forger DB., Francey LJ., Fu Y-H., Gachon F., Gatfield D., de Goede P., Golden SS., Green C., Harer J., Harmer S., Haspel J., Hastings MH., Herzel H., Herzog ED., Hoffmann C., Hong C., Hughey JJ., Hurley JM., de la Iglesia HO., Johnson C., Kay SA., Koike N., Kornacker K., Kramer A., Lamia K., Leise T., Lewis SA., Li J., Li X., Liu AC., Loros JJ., Martino TA., Menet JS., Merrow M., Millar AJ., Mockler T., Naef F., Nagoshi E., Nitabach MN., Olmedo M., Nusinow DA., Ptáček LJ., Rand D., Reddy AB., Robles MS., Roenneberg T., Rosbash M., Ruben MD., Rund SSC., Sancar A., Sassone-Corsi P., Sehgal A., Sherrill-Mix S., Skene DJ., Storch K-F., Takahashi JS., Ueda HR., Wang H., Weitz C., Westermark PO., Wijnen H., Xu Y., Wu G., Yoo S-H., Young M., Zhang EE., Zielinski T., Hogenesch JB. 2017. Guidelines for Genome-Scale Analysis of Biological Rhythms. Journal of biological rhythms 32:380–393.

Kramer PJ. 1936. EFFECT OF VARIATION IN LENGTH OF DAY ON GROWTH AND DORMANCY OF TREES. Plant physiology 11:127–137.

Law CW., Chen Y., Shi W., Smyth GK. 2014. voom: Precision weights unlock linear model analysis tools for RNA-seq read counts. Genome biology 15:R29.

Loader C. 2013. locfit: Local Regression, Likelihood and Density Estimation. Comprehensive R Archive Network (CRAN).

Love MI., Hogenesch JB., Irizarry RA. 2016. Modeling of RNA-seq fragment sequence bias reduces systematic errors in transcript abundance estimation. Nature biotechnology 34:1287–1291.

Love MI., Huber W., Anders S. 2014. Moderated estimation of fold change and dispersion for RNA-seq data with DESeq2. Genome biology 15:550.

Microsoft., Weston S. 2017. foreach: Provides Foreach Looping Construct for R. Comprehensive R Archive Network (CRAN).

Patro R., Duggal G., Love MI., Irizarry RA., Kingsford C. 2017. Salmon provides fast and bias-aware quantification of transcript expression. Nature methods 14:417–419.

Ritchie ME., Phipson B., Wu D., Hu Y., Law CW., Shi W., Smyth GK. 2015. limma powers differential expression analyses for RNA-sequencing and microarray studies. Nucleic acids research 43:e47.

Robinson MD., Smyth GK. 2007. Moderated statistical tests for assessing differences in tag abundance. Bioinformatics 23:2881–2887.

Saito T., Rehmsmeier M. 2017. Precrec: fast and accurate precision-recall and ROC curve calculations in R. Bioinformatics 33:145–147.

Smyth GK. 2004. Linear models and empirical bayes methods for assessing differential expression in microarray experiments. Statistical applications in genetics and molecular biology 3:Article3.

Soneson C., Love MI., Robinson MD. 2015. Differential analyses for RNA-seq: transcript-level estimates improve gene-level inferences. F1000Research 4:1521.

Wu G., Anafi RC., Hughes ME., Kornacker K., Hogenesch JB. 2016. MetaCycle: an integrated R package to evaluate periodicity in large scale data. Bioinformatics 32:3351–3353.

Young MW., Kay SA. 2001. Time zones: a comparative genetics of circadian clocks. Nature reviews. Genetics 2:702–715.

